# Comparison scanning generalized eigendecomposition separates temporal dynamics of alpha oscillations during inhibition of spatialized acoustic distractors

**DOI:** 10.64898/2026.04.23.720427

**Authors:** T.J. Harlow, H. Korsu, L. Almowaly, M. Corcoran, S. Cole, J.J. Chrobak, H.L. Read

## Abstract

Lateralized alpha-band oscillations are thought to reflect distractor inhibition through suppression of cortical regions processing spatially-localized distracting stimuli. Standard lateralization indices (LI) quantify hemispheric asymmetries in spectral power over large time windows, while characterizations of the time-varying dynamics of alpha-mediated distractor inhibition are lacking. Here we evaluate comparison scanning generalized eigendecomposition (csGED), a multivariate signal processing technique, for its efficacy in addressing questions related to alpha-mediated distractor inhibition using a simulation of same-frequency sources at symmetric cortical locations adapted from Zuure and Cohen (2021). We show that while LI accurately captures topographic power asymmetries, csGED is effective at recovering source-projections for bilateral, same-frequency activity across a wide range of signal-to-noise ratios (SNRs). We further extend these models to a pilot sample (N = 11) performing a speech-in-noise task using spatialized naturalistic distractors through individualized head-related transfer functions (HRTFs). Our results demonstrate the efficacy of GED to characterize source projections during spatialized distractors, and provide preliminary evidence for shifts in oscillatory activity in both the alpha (7 - 13 Hz) and beta frequency ranges (15 - 25 Hz) during spatialized speech-in-noise tasks. Together, these results demonstrate the feasibility of csGED for investigating temporal dynamics of lateralized distractor inhibition and motivate larger confirmatory studies.

## Introduction

Selective attention enables individuals to focus on foreground speech while ignoring task-irrelevant background sounds, which is essential for functioning in everyday auditory environments. However, the neural mechanisms underlying this process are not yet fully understood. One proposed explanation is that synchronous alpha oscillations play a key role in distractor inhibition. Specifically, increases in alpha activity are thought to reflect the active suppression of task-irrelevant auditory information, with synchronous alpha oscillations potentially supporting coordinated inhibition across cortical regions (Bonnefond & Jensen, 2017; Jensen & Mazaheri, 2010). Accordingly, alpha activity is thought to increase when people are cued to ignore sounds, while alpha levels decrease when people are cued to attend to upcoming sounds, suggesting a functional role in blocking irrelevant information based on task demands. In lateralized experimental paradigms, this manifests as hemispheric asymmetries in alpha power as quantified by lateralization index (LI; Fiedler et al. 2019). More recently, however, work has specifically incorporated linear modeling approaches to investigate how asymmetries in both periodic and aperiodic activity shift with spatial location (Bender et al., 2025).

Despite these advances in characterizing potential mechanisms for alpha oscillations in distractor inhibition, several open questions remain. First, while lateralization indices summarize asymmetries in alpha power, they do not identify the topographically distinct source-projection patterns which isolate source activity. Consistent with this, it is well known that channel-level analyses are limited in their ability to isolate narrowband activity from sources with similar activity, a problem that is of particular significance when examining bilateral, same-frequency sources. Finally, lateralization indices characterize power asymmetries over discrete time windows, limiting their efficacy in characterizing the time-varying activity patterns of these neural sources over the duration of these tasks. In line with these open questions, multivariate source separation techniques offer a principled framework for addressing these limitations. In particular, generalized eigenvalue decomposition (GED) has been demonstrated to be an effective tool for source separation, band-delimitation, and condition contrasting (Cohen, 2022). Zuure and Cohen introduced comparison scanning GED (csGED), which scans over a range of frequencies to identify those that are maximally separable in their covariance structure between two experimental conditions (Zuure & Cohen, 2021).

In the following work, we evaluate csGED’s efficacy in lateralized source recovery through a novel simulation of same-frequency, alpha-mediated distractor inhibition. We demonstrate that while lateralization indices accurately characterize differences in topographical power distribution between conditions, csGED recovers which sources contribute to those asymmetries, providing complementary but distinct measures of distractor inhibition. Then, we apply csGED on a pilot sample of 11 participants during a speech-in-noise task with variation in spatial background sound distractors. We find preliminary evidence for differential modulation of alpha and beta oscillatory activity in the left-hemisphere when the attended foreground digits play in conjunction with background sounds. While these results should be interpreted with caution given the small sample-size, they demonstrate the efficacy of csGED for condition contrasting in studying temporal dynamics during distractor inhibition tasks and motivate future complementary investigations.

## Methods & Materials

### Participants

All experimental procedures were in accordance with a University of Connecticut Institutional Review Board (IRB) approved protocol. All participants were provided with a verbal and written description of the procedures and rationale, and all participants provided informed consent. Subjects received payment or course credits for participation. An otoscopic inspection confirmed the absence of earwax buildup. Participants reported no neurological disorders or use of prescription medications. The initial data collection resulted in 13 healthy young adults. Subsequent exclusion criteria was based on two factors: (i) excessive EEG artifacts amounting to greater than two corrupted recording blocks, and (ii) behavioral performance below the 1st percentile in foreground sound identification across all sound conditions, performance levels that did not differ significantly from chance, indicating a lack of task engagement or comprehension. Thus, this study includes data from a total of 11 subjects. Demographic information for one participant was unavailable due to corruption of their questionnaire data file. Of the 10 participants with complete demographic information, there were 5 females with a mean age of 20 (+2.9) years old and 5 males with a mean age of 23.8 (+4.8) years old. The total population age mean and standard deviation was 21.9 (+4.2) years old.

### Experimental paradigm and data collection

All experimental procedures were conducted in the shared EEG laboratory at the University of Connecticut’s Institute for Brain and Cognitive Sciences (IBACS), inside an acoustically shielded chamber manufactured by Industrial Acoustics Inc. Auditory stimuli were presented binaurally through ultra-shielded Etymotic earphones (Intelligent Hearing Systems) at a fixed intensity level that was identical across all participants. Task presentation was controlled in MATLAB (R2024b, The MathWorks) using the Psychtoolbox 3.20 extension (Brainard, 1997). Throughout each trial, participants maintained visual fixation on a cross displayed on a computer monitor positioned at eye level. Each participant completed the experiment across two recording sessions, scheduled on separate days, with each session lasting approximately two hours. At the beginning of each session, a two-minute resting baseline EEG segment was recorded while participants kept their eyes closed. Following this baseline period, the speech-in-noise task was administered in two recording blocks per session, each lasting approximately 20 minutes and separated by a brief break to minimize potential time-on-task effects. Across the two recording days, a total of 600 trials were obtained from each participant. Trials were distributed evenly across background sound categories, yielding 100 trials per category overall. On each trial, a unique time segment of each background sound was played to reduce memorization of the background sounds. Within each session, trials were organized into randomized blocks of 150 trials. Participants performed the standard speech-in-noise task where they ignored the background sounds and attended to and remembered a three digit sequence of numbers (Spatialized Speech-in-noise Task).

### Spatialized Speech-in-Noise Task

After an initial short task to confirm that participants could localize left and right spatial locations (described below), participants were instructed to disregard the background sounds and to report a sequence of three foreground digits after the sound ended by entering their responses on a keyboard following each trial. The background sound was first presented alone for a minimum of 700 ms prior to the onset of the jittered foreground digit sequence. The foreground digit sequence played in combination with the background sounds in the 700 to 3,000 ms time period. The total duration of each speech-in-noise stimulus was 3.5 seconds, with sound onsets and offsets smoothed using a b-spline filter. Six categories of background sounds were used: (1) eight-speaker Babble, (2) Factory, (3) Freeway, (4) Running Water, (5) Crackling Fire, and (6) White Noise. Distinct segments from each background category were used on every trial to reduce the likelihood of memorization, and foreground stimuli consisted of digits zero through nine spoken by eight different male talkers from the TI46 LDC Corpus (Liberman, 1991), consistent with a prior speech-in-noise behavioral study (Clonan et al., 2025). Both the background sound category and digit sequences were randomly assigned on each trial.

Background sounds were spatialized through an individualization of the standard head-related transfer functions (HRTFs) implemented in MATLAB. Specifically, a KU100 head model taken from the Sadie II database served as the basis for the HRTF ([Sadie II Project](https://www.york.ac.uk/sadie-project/database.html)). First, each individual participants’ head perimeter and intertragus distance are measured, from which each participants desired objective azimuth can be calculated directly from the polynomial fits to KU100 headmodels provided by Gutierrez-Parera and colleagues 2022 (Gutierrez-Parera et al., 2022). During the initial block of 150 trials, participants performed a spatial localization task to identify the left versus right spatial location of the background sounds that were presented in random order. This task confirmed that participants perceived left versus right spatial locations for each background sound (**Results**).

Foreground digit onsets were jittered across trials for each digit within a sequence. The first digits played during a time window of 700 to 1100 ms with a mean and standard deviation of 896.6 (+ 119). The second digits played during a time window of 1060 to 2010 ms with a mean and standard deviation of 1524 (+182). The third digits played during a time window of 1510 to 2990 ms with a mean and standard deviation of 2152 (+ 224). This minimum 700 ms delay for the first digit was chosen in order to optimize the potential for neural and behavioral classification of the background sounds, as we demonstrated previously (Zhai et al., 2020). Moreover, by starting the attended digit sounds after 700 milliseconds, we aimed to minimize the suppression of the attended speech-evoked response potentials, as demonstrated previously (Papesh et al., 2015). Foreground sounds were 3 decibels (3 dB) higher sound-level than the background sounds with a signal-to-noise ratio (SNR) of +3 dB, in order to maximize the foreground speech-evoked response amplitudes and the task performance, as shown previously (Kim et al., 2021; Muncke et al., 2022).

### EEG acquisition and processing

EEG recordings were collected using a 64-channel electrode cap arranged according to the standard 10–20 system (Brain Vision system). Signals were recorded online with PyCorder software at a sampling rate of 5 kHz, and electrode impedances were maintained below 10 kΩ throughout acquisition. All subsequent preprocessing and analysis steps were performed offline in MATLAB (R2024b). During preprocessing, EEG signals were first re-referenced to a common average reference. The data were then downsampled to 250 Hz after applying an anti-aliasing filter implemented through the EEGLAB MATLAB toolbox. Independent Component Analysis (ICA) was performed following standard EEGLAB-recommended procedures. To optimize component separation, the data were initially passed through a narrow bandpass filter spanning 1–30 Hz prior to decomposition. Independent components were computed using the logistic infomax ICA algorithm (Bell and Sejnowski, 1995). The resulting components were subsequently transferred to data that had been more broadly filtered between 0.1 and 50 Hz to support artifact rejection. Component classification relied on both automated and manual evaluation: ICLabel (Pion-Tonachini et al., 2019) was used to generate preliminary labels, and components were then visually inspected based on their scalp topographies and temporal activity patterns. Components associated with ocular, muscular, and cardiac artifacts were identified through this process and removed before further analysis.

### SIMULATION analyses

For simulated analyses, we use a cortical surface model composed of 2004 dipole locations, each of which associates with a projection weighting matrix taken from (Cohen 2020). Similar to prior work developing the comparison scanning procedure, we consider two contrasting conditions. Previously, these sources differed in both frequency and location (Zuure and Cohen 2021), but here we restrict our analyses to two conditions with differing alpha-band source locations. Both conditions have a center frequency of 10 Hz, and are wrapped with a Gaussian envelope with a variance of 500ms. The relative SNR between source amplitudes on any given trial is varied between 0.5 and 5 in steps of 0.5. The amplitudes of left/right alpha are then calculated with ρ = 10 · *snr*, following which the dominant amplitude is taken as 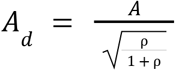 and the corresponding nondominant source then being 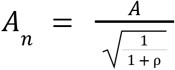. For simplicity, we fix the total amplitude *A* to be three, and maintain a consistent source of brownian noise generated at each leadfield for both conditions. Scalp projections are then estimated through the product of the leadfield and the activity for each corresponding dipole, and ground truth projections being based on the absence of any noise component. The above parameters result in 125 unique left and right trials at each given SNR level.

### Comparison scanning

Comparison scanning generalized eigendecomposition (csGED) is implemented according to Zuure and Cohen (2021), and adapted here for contrasting conditions with left and right lateralized distractor conditions at each scanning frequency of interest for both simulated and empirical datasets. For csGED, we scanned 30 frequencies logarithmically spaced between 3 and 30 Hz using complex Morlet wavelet transform:

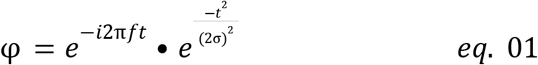

Here, “*i”* represents the complex variable, “*f”* is the center frequency, and “*σ” is* the time bandwidth of the wavelet. Cycles (*c*) were linearly spaced from 3 to 12 with each center frequency, with the time bandwidth 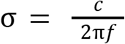. The real part of the resulting narrowband oscillatory signal is then used for per-trial covariance channel-by-channel covariance estimation at a given frequencies. For simulated data, covariance matrices are estimated using a 750ms window about time zero. For empirical data, covariance matrices are estimated from both pre and post foreground stimulus onset, and subsequently separated into left and right lateralized distractor conditions before averaging. We additionally reject outliers on the basis of a z-score threshold of |z| > 2.721 (99.7 %) relative to the mean covariance within each condition x frequency set (Cohen 2020). After which condition means are recomputed before performing generalized eigendecomposition (GED). For GED, two versions are performed: once for maximizing contrast between left-trials and regularized right-trial covariances, and vice versa. Following Zuure and Cohen (2021), regularization is done using ridge regularization, with the corresponding MATLAB notation being:

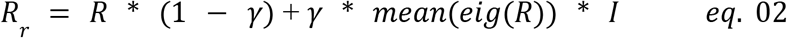

Where *R* represents the reference matrix for a particular GED and *R*_*r*_ then is the regularized reference matrix using a γ of 0.01. Consistent with prior literature, only the leading eigenvector with the largest corresponding absolute eigenvalue is taken for each frequency x condition combination before normalization to unit length. This results in a spectrum of eigenvalues reflecting the degree of separability of simulated and empirical conditions at each frequency. For each of which the corresponding forward model can be computed simply through the product of the eigenvector and the signal covariance matrix (Cohen 2020).

Group eigenvalue spectra are represented similarly to that of Zuure and Cohen (2021). Specifically, for each participant the eigenvalue spectrum findpeaks is used to identify at most four peaks with a minimum peak distance of 0.075 in log10-based frequency. This roughly corresponds to a 2.6 Hz wraparound at a 10 Hz center frequency. From each participants’ identified peaks, kernel density estimation (KDE) is used to identify smoothed spectra using gaussian kernels with a bandwidth of 0.05 in log10-based frequency. Peaks in the group averaged, kernel-smoothed spectra were used to define frequency bands of interest following the golden-ratio rule of 2.33 Hz ratio between center frequency and bandwidth (Klimesch 2018).

## Multitaper analyses

Multitaper analyses are conducted in two general forms: (1) for the estimation of channel-level LIs during simulated analyses, and (2) using a multitaper spectrogram approach for empirical comparisons of oscillatory dynamics. For LI, power spectral density is first estimated for each individual channel using Thomson’s multitaper method (Thomson 1982). Specifically, we use a time half-bandwidth product of two and three tapers, evaluated at frequencies ranging from 3 to 30 Hz. Multitaper analyses were conducted over the same analytic window as csGED approaches, that being a 750ms window wrapped around time zero. Aperiodic activity is removed using a linear fit applied in log-log space over the range of 2 to 40 Hz, from which periodic activity is expressed as decibels (dB). Alpha power is that taken as the cumulative power above zero within the range of 7 - 13 Hz, with LI between left and right conditions subsequently being estimated using:

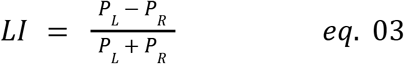

with *P*_*L*_ and *P*_*R*_ corresponding to the cumulative power for a given channel during left and right dominant trials, respectively.

Time-frequency representations of empirical EEG data were produced using sliding-window multitaper spectrograms (Chronux toolbox; Mitra & Bokil, 2007). Spectrograms were made on a per-trial basis with a 400ms sliding window, overlap of 15ms, a time half bandwidth of 1.5 with two tapers, over the frequency range of 2 to 50, and zero-padding was performed using a factor of two. In order to isolate time-varying periodic activity, multitaper spectrograms were parametrized using an aperiodic model including a knee parameter:

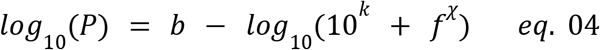

where *b* is the model slope intercept, *k* the frequency knee parameter, and *χ* being the spectral aperiodic. Model parameters were estimated using via nonlinear least squares (*lsqnonlin*) with the following bounds: *b* ϵ [-20 20], *k* ϵ [-8 8], *χ* ϵ [0.2 6.0]. Following identification of the aperiodic fit, spectrograms were detrended through subtraction in log space and expressed in decibels. Detrended multitaper spectrograms were centered to each trial’s time-averaged mean. Subject-specific component eigenvectors identified from group-averaged peaks in csGED separability spectra were then used to spatially filter single-trial multitaper spectrograms by projection through the corresponding component weights.

### Statistical analyses

For simulated comparisons of csGED and LI, bootstrap resampling with replacement is performed using 250 iterations per SNR condition. Ground truth lateralization maps were computed through the channel-wise dB ratio of mean squared amplitude between left and right dominant trials without the inclusion of the noise component. For csGED, contrast maps were derived using a normalized contrast between the left and right forward models (*aL, aR*, respectively):

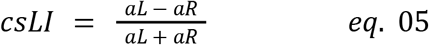

Whereas for LI, the standard lateralization index was computed from detrended alpha power in the range of 7 to 13 Hz. Statistical comparisons of contrast maps were performed using the squared Pearson correlation between each method and those resulting from ground-truth projections.

The accuracy of source recovery was measured using cosine error (Nikulin et al., 2011), as defined by:

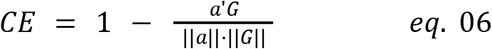

where *a* reflects the estimated forward model and *G* being the true dipole projection through the leadfield. For *CE*, values of zero indicate perfect recovery while values of one indicate complete orthogonality. The *CE* measure is also scale and sign invariant, and as such was computed independently for left and right trials before averaging.

Given our limited sample size (N = 11), there are concerns about the internal validity of csGED components over conventional inferential statistics. Here, we address this concern by investigating the consistency of forward component maps with *R*^2^, a procedure consistent with the gedBounds procedure developed by Cohen (2021). We find high within-frequency consistency, indicated by large correlations in topographic maps by adjacent frequencies, and lower correspondence between distant frequencies. Similarly, at the group level, consistency of peak locations in the eigenvalues of separability spectra is assessed through our kernel density estimated spectra as described above.

Finally, differences in GED-projected spectrograms between left and right distractor conditions were assessed using nonparametric cluster-based permutation testing (Maris & Oostenveld, 2007). At each time-frequency bin, a paired test statistic was computed across subjects comparing left-vs. right-distractor trials projected through the left-component eigenvector. Adjacent bins exceeding a bin-level threshold (p < 0.05, uncorrected) were aggregated into clusters by summing their test statistics. The cluster-level significance was determined by comparing observed cluster masses to a null distribution generated from 1000 random permutations of the condition labels. Clusters exceeding the 95th percentile of the null distribution were considered significant (Maris & Oostenveld, 2007).

## Data and code availability

Simulated data, code, and derivatives from real data are available at the following link: https://github.com/TylorJHarlow/Comparison-scanning-generalized-eigendecomposition-separates-temporal-dynamics-of-alpha-oscillations

## Results

### Simulated Hemispheric Spatialization of Alpha Oscillations

To validate the efficacy of GED to capture neural sources associated with distracting background sounds, we simulate a dataset that models bilateral alpha oscillations during performance of a speech-in-noise task in the time period when there was only background sound and no target foreground speech digit sequence (**Methods**). The simulated data are based on those developed by Cohen and colleagues (**Methods;** Cohen 2020; Zuure and Cohen 2021; Cohen 2022). Neural activity is modeled using two principal sources through a standard leadfield matrix. This includes aperiodic activity with an exponent of 2, generated across all 2004 dipole sources. Alpha oscillations are simulated as higher in the left brain hemisphere when the ignored background sounds are playing on the right side of the head (**Fig. 1A**). Conversely, alpha oscillations are simulated as higher in the right brain hemisphere when the ignored background sounds play on the left side of the head. These simulated alpha oscillations have a center frequency of 10Hz, and are tapered using a Gaussian envelope with a variance of 500 milliseconds. The simulated higher power alpha oscillations in the left brain hemisphere elicited by background sounds played from the right spatial location are illustrated (**Fig. 1A**, blue line).

**Figure 1.**
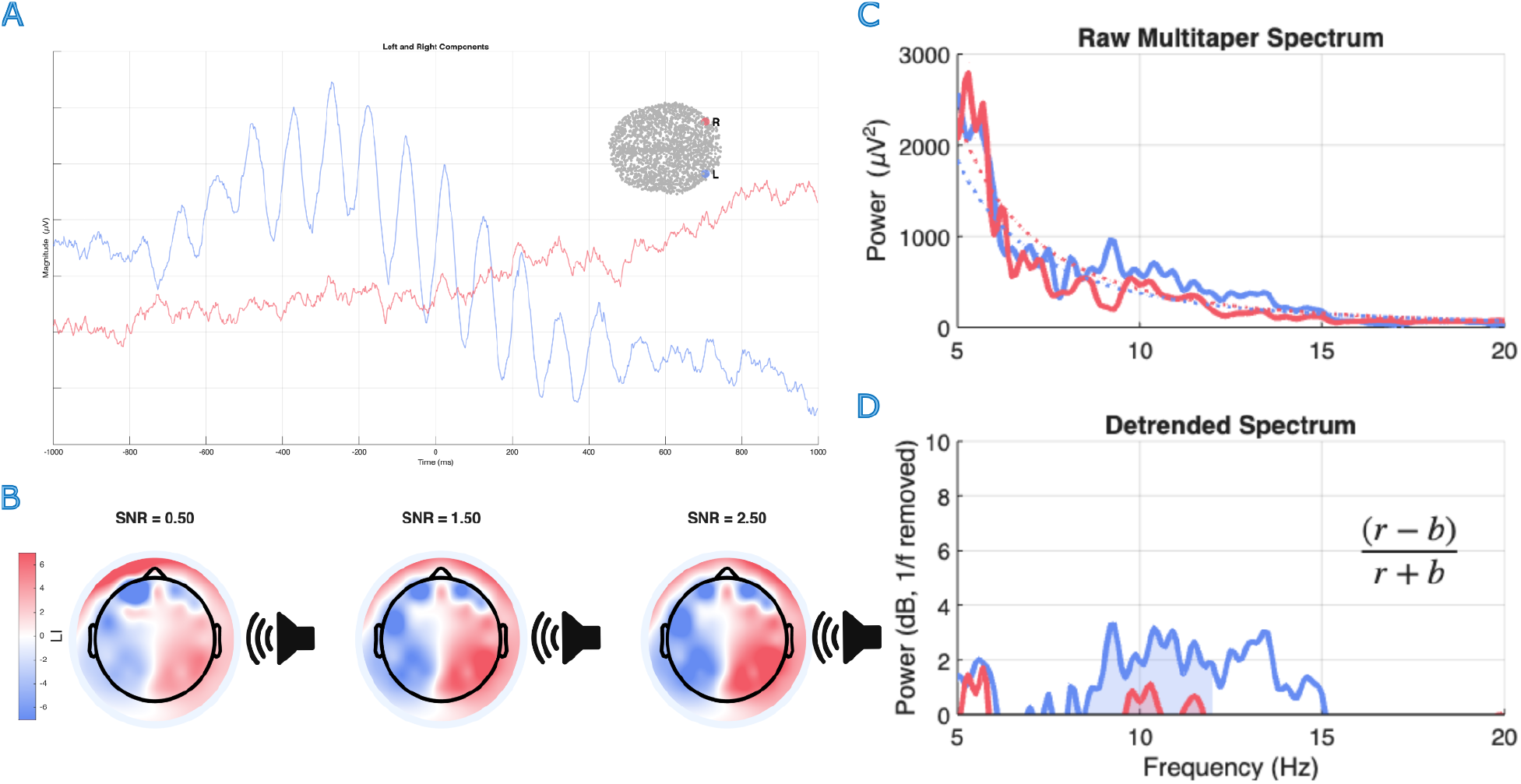
Alpha-mediated distractor inhibition: A) Provides a visual representation of aperiodic and alpha components on a single trial with and ignored sound played on the right spatial location, and alpha highest in the left brain hemisphere. The left (blue line) and right (pink line) alpha oscillation source locations are illustrated topographically within the leadfield model icon shown (Methods). B) Ground truth topographic lateralization measures generated in the absence of noise for three example hemispheric signal-to-noise ratios (SNRs). Colorbar ranges from −7 to +7 for lateralization index (LI), with blue indicating higher alpha in the left brain hemisphere and red indicating higher alpha power on the right brain hemisphere. C) Standard multitaper spectrogram for a single example trial with left-lateralized alpha as in **A.** D) The corresponding aperiodic component (1/f) detrended multitaper spectra, with shaded AUC for estimation of the lateralization index (LI).

Simulated trials also varied in the degree of lateralization of alpha power generated from the two given posterior electrode sources through a mutual alpha power level hemispheric signal-to-noise ratio (SNR) parameter. This SNR parameter controlled the relative distribution of a fixed amount of alpha power between the two hemispheric alpha source locations. Our GED simulation demonstrates the ground truth topographic projection of alpha power across three simulated versions of interhemispheric SNR variations for alpha (**Fig. 1B**). Characterizing the different levels of hemispheric lateralization of alpha power has generally relied on what is referred to as the Lateralization Index (LI), which consists of the ratio between electrode cluster power differences between conditions. Here, we simulate the aperiodic and periodic alpha components as shown in the single-trial example power spectrum plot (**Fig. 1C**). An aperiodic-detrended version of this analysis is also shown (**Fig. 1D)**, for the right versus left hemispheric channels. Such power analyses are typically used to compare changes in alpha across brain hemispheres when ignored sounds are played from differing spatial locations.

We next compare the ability of the lateralization index versus that of the comparison scanning generalized eigenvector decomposition (csGED) to recover ground truth lateralization patterns. Here, we simulate increased alpha power levels contralateral to the ignored sounds on the left versus right side of the head (**Fig. 2A**). Prior studies have demonstrated how csGED quantifies the frequencies for maximally separating two different cognitive task conditions and allows one to identify brain sources contributing to particular frequencies (Zuure and Cohen 2021). On each trial, the alpha power is simulated to be maximal in the central-parietal regions contralateral to the ignored sound (**Fig. 2A**). Trials are narrowband filtered using Morlet wavelets (**Methods**), sorted by condition, and used to construct per-trial covariance matrices. An example condition-averaged covariance matrix is shown for alpha left-dominant trials simulating ignored background sound spatially located on the right side of the head (**Fig. 2B)**. Following the comparison scanning csGED between left and right trial covariances, the leading Eigenvector is evaluated on two complementary measures. First, for its ability to recapitulate topographic power differences (**Fig. 2C**, R^2^), and second to quantify the correspondence between hemispheric alpha SNR and cosine error projections (**Fig. 2D**). Unsurprisingly, we find that the lateralization index is effective at capturing lateralization in power between left and right distractor conditions (**Fig. 2C**, blue line simulation). We also find that the lateralization index (R^2^) significantly accounts for the variance and improves as the inter-hemispheric alpha SNR increases from 0.5 to 5 (**Fig. 2C**, blue line, β_*MT*_ = 0.01, p = 4.81e-18). In contrast, the comparison scanning csGED lateralization index (R^2^) varies less with changes in the inter-hemispheric alpha SNR (**Fig. 2C**, β_*csGED*_ = 0.002, p = 2.69e-6). Hence, csGED is markedly less effective at characterizing the lateralized power asymmetries. In contrast, we find that csGED cosine error is smaller and the mean value decreases systematically with the increase in inter-hemispheric alpha SNR (SNR: 0.5 to 5), consistent with recovery of ground truth source topographies (**Fig. 2D**, pink line, β_*GED*_ = -0.002, p = 3.14e-111). In contrast, the multitaper lateralization index (cosine error) does not vary significantly or systematically with the hemispheric alpha SNR (**Fig. 2D**, blue line β _*MT*_ = -0.0009, p = 0.566). These results demonstrate the efficacy of the standard lateralization index (R^2^) to describe where and how alpha power differs between two simulated conditions. Moreover, the csGED accurately quantifies the change in cosine error with increasing alpha power SNR differences across brain hemispheres. Thus, these two metrics provide distinct, complementary approaches to characterizing changes in alpha oscillatory dynamics when background sounds vary in spatial location.

**Figure 2.**
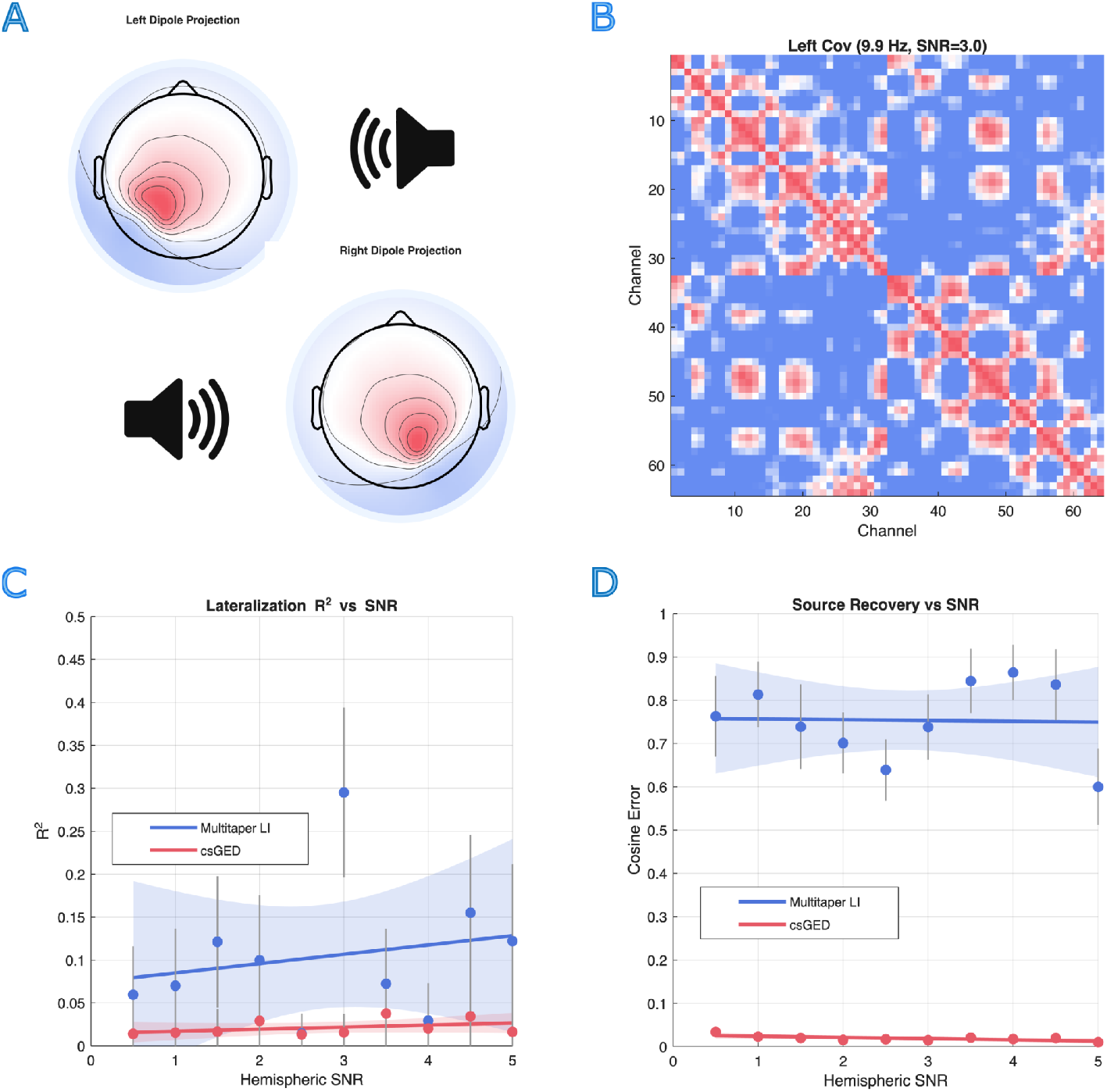
GED captures alpha lateralization across SNRs: A) True topography projections for left and right alpha source activity. B) Visualization of example alpha-band channel covariance matrix built from left trials. C) Comparison of R^2^ of GED and LI maps to ground truth alpha power lateralizations as a function of SNR. Shaded regions denote 95% confidence intervals for fit lines. Grey bars denote standard errors in mean R^2^across permutations. D) Same as **Figure 2C** but using cosine error metric for capturing distribution of source activity.

### Alpha Lateralization During Speech-in-Noise Task Performance

Our speech-in-noise task consists of four major recording blocks, each consisting of 150 trials, conducted over two days. For the first block, participants are required to confirm the perceived direction of the individually spatialized background sounds on each trial in order to confirm that the individualized HRTFs are correct (**Methods**). During this recording block, participants correctly identify the background distractor sound location with a performance accuracy of 91.03 % (SEM + 5.07%). This indicates that the individualized HRTFs produce a reliable perception of lateralization across six different background sound categories. In experimental blocks 2-4, participants accurately identified foreground target digits 99.3% (SEM + 0.0024%), consistent with the high speech-in-noise task signal-to-noise ratio (SNR = + 3 dB) used in this pilot study.

Next, given our limited sample size (N = 11), we address the internal validity of csGED components before secondary analyses based on separability spectra. Specifically, we evaluate the consistency in pairwise correlations between eigenvector topographies across all scanning frequencies separately for left and right csGEDs during the background sound-only time period before the target foreground digit sequence onset (**Fig. 3A, B**). We implemented the gedBounds procedure of an individual participant basis (**Methods**) using eigenvector correlation matrices according to prior work. This procedure involves the iterative application of *dbscan* over various clustering sensitivity levels (Cohen 2021). We subsequently and then averaged across subjects to indicate the consistency of cluster identification across participants. Both conditions present a highly diagonal structure, indicating adjacent frequencies having predominantly similar topographic forward model projections, and that this is common across participants. In contrast to this, distant frequency components tend to have lower correlations, indicating that they represent spatially distinct auditory inputs. In line with our separability spectra, we find multiple significant clusters, providing preliminary evidence for multiple shifts in alpha oscillatory activity related to the background distractor sounds (**Fig. 3A, B**). This indicates that csGED identifies spatially distinct sources at a variety of frequency bands, rather than recovering a single global spatial pattern, as is the case for standard lateralization indices. These findings are consistent with the band-delimitation properties of GED described in prior work (Cohen, 2021).

**Figure 3.**
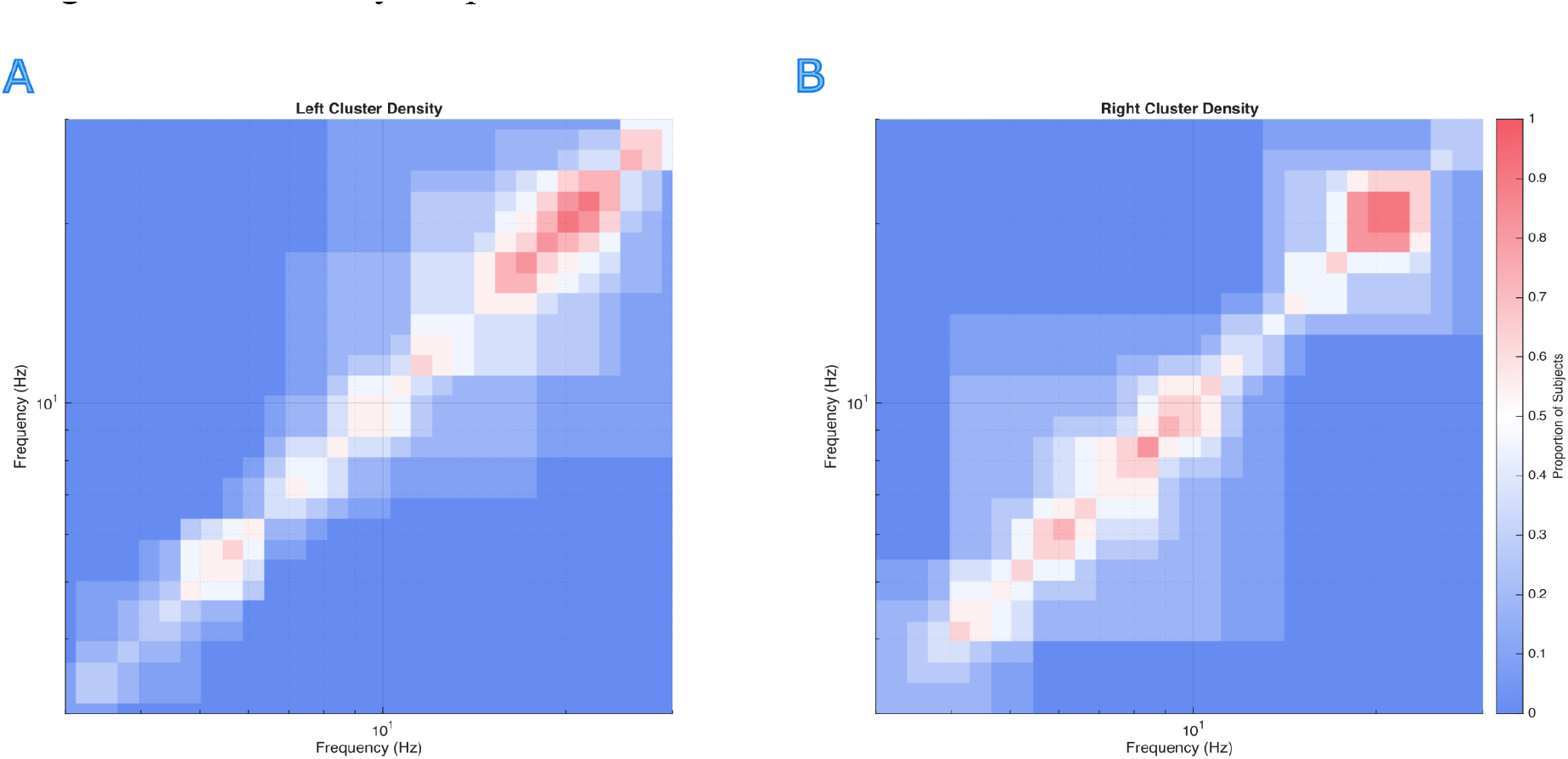
Specificity and generalizability of component peaks. A) Matrices reflecting consistency of clusters in topographic correspondence (R^2^) across subjects for blocks 2-4, during the ignored background sound only time period (0 - 700ms) for left spatial distractors. B) Same as **A**, but for background sounds spatially positioned to the right side of the head. For both A and B, the colorbar ranges from 0 to 1, reflecting consistencies across subjects in frequency-specific topographies via the R^2^.

### Time-varying oscillatory dynamics

Having established the internal validity of csGED components, we next look to evaluate the spectral separability during blocks 2 - 4 of our experiment. The group-level, kernel density estimation (KDE) smoothed Eigenvalue for the broadband spectra are shown for the background only time window (**Fig. 4 A, B**, “pre-foreground”, light red & blue) and for the combined foreground-background sound time window (**Fig. 4 A, B**, “post” foreground onset, dark red & blue). For left background sound spatial location, multiple peaks are evident in the spectra with prominent peaks appearing in the alpha (7 - 13 Hz) and beta (15 - 30 Hz) frequency ranges when the background sound plays in isolation before the foreground attended digit sequence onsets (**Fig. 4 A, B**, light lines). When the background sound is located to the left, the alpha peak frequency is reduced in amplitude following foreground stimulus onset (**Fig. 4A**, dark red line, 10 Hz). A similar decrease in alpha peak frequency at 10-12 Hz is observed when the background sound is located to the right and the foreground sound onsets (**Fig. 4B**, dark blue line, 10-12 Hz). Together, this indicates that oscillatory signals corresponding to distractor inhibition are modulated by target stimulus onset for both spatial locations of the background sound.

**Figure 4.**
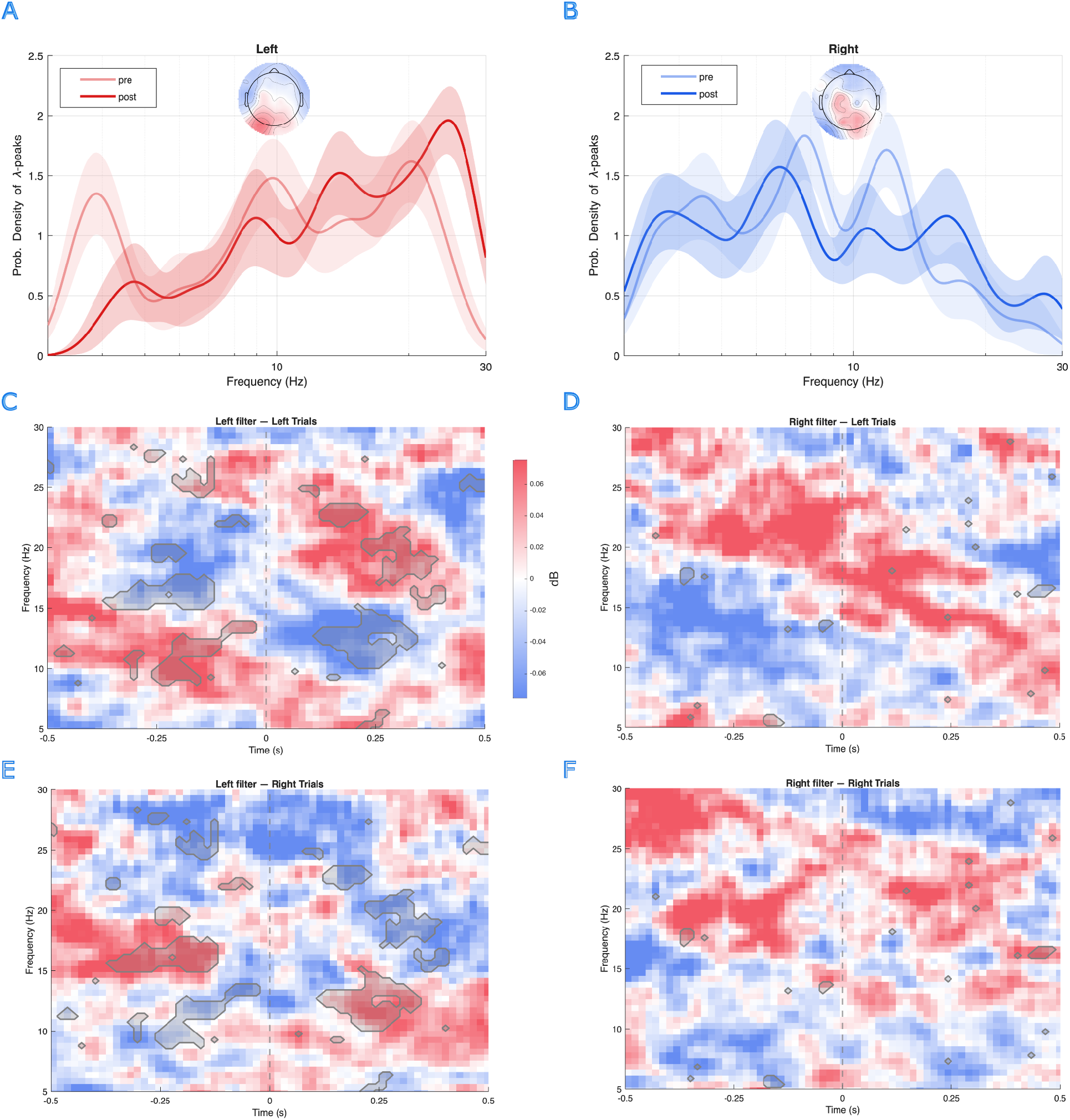
Comparison scanning spectra and time series representations: A) Left trial separability spectra derived using comparisons scanning during pre and post foreground onset periods during blocks 2-4. Topographic plot shows high alpha power in parietal-occipital areas of the left brain hemisphere. B) Same as **Figure 4A** but for trials where the background sound was located on the right side of the head. Topographic plot shows source projections at group-averaged alpha frequency. C) Multitaper spectrograms for left trials from blocks 2-4 through the left component eigenvector. E) Same as **Figure 4C** but projection is onto right trials. D) Multitaper spectrograms for left trials from blocks 2-4 through the right component eigenvector. E) Same as **Figure 4D** but projection is onto right trials. C-F) Gray shaded regions denote significant differences in activity levels between left and right trials, as determined through nonparametric permutation testing.

In order to visualize these changes over time, we plot the csGED-projected detrended multitaper spectrograms relative to the foreground attended digit onset (**Fig. 4 C, D** t=0) for both left (**Fig. 4C**) and right (**Fig. 4D**) spatially located background sound trials. Shaded regions indicate significant clusters, as determined through a two-tailed nonparametric permutation test, where left-dominant trial activity significantly differs from that of right-dominant trials (**Fig. 4C vs D**, p < 0.05; Maris & Oostenveld, 2007). Opposing alpha and beta dynamics are notable between the left (**Fig. 4C**) versus right (**Fig. 4E**) dominant hemispheric clusters which correspond to the background sound localized to the right and left side of the head, respectively. Alpha activity is high during the background sound-only time period (t=-0.5 to 0 seconds), but rapidly desynchronizes, decreasing in power, with the attended foreground speech digit onset (t = 0 seconds) for trials where the background sound is spatially located on the left side of the head (**Fig. 4C**). In contrast, alpha oscillations in the left-hemispheric cluster elicited with background sound spatially located on the right side of the head (**Fig. 4E**) have increased synchronization with the foreground digit onset (**Fig. 4E**, following t = 0 s). Beta oscillatory activity demonstrates the opposing trend, where it synchronizes with foreground stimulus onset during left trials (**Fig. 4C**, 15-30 Hz, and vice versa for the right-component trials during left distractor trials (**Fig. 4D**, 15-30 Hz). The elevated alpha activity we observe during the background sound-only period (−500 to 0 ms) contralateral to the distractor, followed by desynchronization at foreground onset, parallels the anticipatory alpha lateralization reported by Wöstmann et al. (2019), who found that contralateral alpha increases prior to stimulus onset reflects proactive distractor suppression.

The above results should be interpreted with caution, as the corresponding analysis for the right-component Eigenvector did not reveal comparably strong opposing alpha-beta dynamics. While some evidence suggests elevated alpha power during left component trials during the post-foreground target onset, alpha and beta modulation was less consistent across participants (**Fig. 4D**,**F**). Given the limited sample size, such incongruencies could reflect low statistical power, or possible left-hemispheric dominance of speech-processing. In either case, the above results overall highlight the efficacy of csGED for investigating oscillatory activity when listening to foreground sounds combined with spatial distractor background sounds, demonstrate its applicability to empirical data, and highlight the need for future investigations with larger experimental cohorts.

## Discussion

The present study evaluated the utility of comparison scanning generalized eigendecomposition (csGED) for characterizing spatially lateralized oscillatory activity engaged during performance of a speech-in-noise task. Using both simulated datasets and empirical EEG recordings obtained during a spatialized speech-in-noise task, we demonstrate that csGED provides improved recovery of source-separated activity compared to traditional lateralization index (LI) approaches and reveals temporally dynamic oscillatory patterns associated with distractor processing. Together, these findings extend prior work on alpha-mediated inhibition of task-irrelevant sensory input by demonstrating that multivariate decomposition approaches can resolve bilateral same-frequency sources and capture time-varying oscillatory dynamics that are not accessible through conventional channel-level metrics.

### Simulation validates csGED for source recovery

In the simulated analyses, we found that the LI reliably captured hemispheric differences in power distribution, but csGED more accurately recovered the spatial projections corresponding to the underlying sources across a wide range of SNR conditions. This distinction highlights an important conceptual difference between the two approaches. While the LI reflects differences in power magnitude across channels, csGED isolates the spatial weighting patterns that defines the contributing neural sources. These findings align with prior work demonstrating the utility of generalized eigenvalue decomposition (GED) for isolating physiologically meaningful components from multichannel electrophysiological recordings (Cohen, 2022). The comparison scanning GED-based methods have previously been shown to improve signal-to-noise ratio and enhance spatial specificity when separating oscillatory sources (Zuure & Cohen, 2021). The present findings extend those results by demonstrating that csGED can be applied specifically to scenarios involving bilateral same-frequency sources, which represent a known challenge for channel-level spectral analyses. Importantly, our simulation results indicate that csGED performance remains robust across increasing SNR levels, suggesting that the method is appropriate for identifying oscillatory sources even under moderate noise conditions.

### csGED in pilot data

In the empirical portion of the study, csGED revealed multiple peaks in the separability spectra, particularly within the alpha (7–13 Hz) and beta (15–30 Hz) frequency ranges. These peaks were observed when the background sound played in isolation (aka, “pre-target onset”) versus when the attended speech digits played together with background sounds (aka, “post-target onset”). Furthermore, the spectral peak probabilities shifted toward lower frequencies following the attended foreground digit onsets. This suggests a modulation of oscillatory activity associated with transitions from distractor suppression to target processing. These findings are consistent with established models of alpha-band activity as a mechanism for functional inhibition (Haegens et al., 2015). In particular, increases in alpha activity have been widely interpreted as reflecting suppression of task-irrelevant information (Jensen & Mazaheri, 2010; Bonnefond & Jensen, 2017). In lateralized attention paradigms, alpha power typically increases contralateral to ignored stimuli and decreases contralateral to attended stimuli, often thought to reflect selective gating of sensory processing (Wöstmann et al., 2019). The opposing alpha dynamics we observe through the left-component Eigenvector for left versus right distractor trials, where alpha synchronizes contralateral to the distractor and desynchronizes ipsilaterally, mirrors the sign-reversed lateralization pattern reported by Wöstmann et al. (2019), who found that alpha power increases contralateral to anticipated distractors and decreases ipsilaterally, independent of target location.

### Limitations and future directions

Several limitations should be considered when interpreting the present findings. First, the empirical dataset consisted of a small pilot sample (N = 11), which limits statistical power and may reduce the generalizability of the observed effects. For example, the weaker beta modulation observed in right-hemispheric analyses may reflect insufficient sample size rather than true hemispheric asymmetry. Second, while csGED provides improved spatial separation relative to channel-level analyses, it does not directly localize neural sources in anatomical space. Future studies combining csGED with source localization methods such as beamforming or inverse modeling could further enhance the interpretability of the spatial components. Third, the present simulations were based on simplified models of bilateral alpha activity. Although these models were designed to approximate realistic neural signals, real neural dynamics may involve more complex interactions between multiple sources and frequency bands. Expanding simulation frameworks to include additional noise sources and temporal variability would provide a more comprehensive evaluation of csGED performance. Similarly, these same simulations could be used for evaluations of recent advancements in modeling of neural time series that have been drawn from dynamical systems research (Brunton 2016; Marrouch 2021).

In addition, future work should extend these analyses to larger experimental cohorts in order to validate the observed oscillatory patterns and improve statistical reliability. Increasing the number of participants would also allow investigation of individual differences in distractor suppression strategies, which may be reflected in variability of component topographies or temporal dynamics. Additionally, future studies could incorporate more complex spatial configurations, such as multi-directional distractor fields or dynamically shifting sound sources. Such designs would enable evaluation of how oscillatory networks adapt to changing spatial environments. Finally, combining csGED with behavioral performance metrics would allow investigation of how oscillatory dynamics relate directly to perceptual outcomes. Establishing such relationships would provide stronger evidence linking neural oscillations to functional mechanisms of selective attention.

## Conclusion

Here, we validate the efficacy of csGED to recover source-projections of same-frequency oscillations under conditions often assumed during inhibition of spatialized distractors. While traditional lateralization indices accurately characterize differences in power between conditions varying in hemispheric dominance, csGED accurately characterizes source-projections, affording investigations into temporally-resolved, source-projection dynamics. This technique was then applied to a pilot sample (N = 11) undergoing a speech-in-noise task with naturalistic background sound distractors. Preliminary results demonstrated multiple shifts in oscillatory activity between left and right spatial locations of ignored background sounds, and opposing alpha and beta activity dynamics. In future work, we look to extend these results to larger sample sizes undergoing similar speech-in-noise conditions at a more challenging SNR that is known to impact behavioral performance levels (Clonan et al., 2025). With this design, we expect to be able to relate temporal properties of distractor and target associated oscillatory activity more directly with measures of behavioral performance.

